# Mindfulness Improves Brain Computer Interface Performance by Increasing Control over Neural Activity in the Alpha Band

**DOI:** 10.1101/2020.04.13.039081

**Authors:** James R. Stieger, Stephen Engel, Haiteng Jiang, Christopher C. Cline, Mary Jo Kreitzer, Bin He

## Abstract

Brain-computer interfaces (BCIs) are promising tools for assisting patients with paralysis, but suffer from long training times and variable user proficiency. Mind-body awareness training (MBAT) can improve BCI learning, but how it does so remains unknown. Here we show that MBAT allows participants to learn to volitionally increase alpha band neural activity during BCI tasks that incorporate intentional rest. We trained individuals in mindfulness-based stress reduction (MBSR; a standardized MBAT intervention) and compared performance and brain activity before and after training between randomly assigned trained and untrained control groups. The MBAT group showed reliably faster learning of BCI than the control group throughout training. Alpha-band activity in EEG signals, recorded in the volitional resting state during task performance, showed a parallel increase over sessions, and predicted final BCI performance. The level of alpha-band activity during the intentional resting state correlated reliably with individuals’ mindfulness practice as well as performance on a sustained attention task. Collectively, these results show that MBAT modifies a specific neural signal used by BCI. MBAT, by increasing patients’ control over their brain activity during rest, may increase the effectiveness of BCI in the large population who could benefit from alternatives to direct motor control.

## Introduction

Through noninvasively decoding user intent in real time, brain-computer interfaces (BCIs) bypass traditional neuromuscular pathways to provide alternative routes of communication and action for those afflicted by paralysis resulting from stroke, trauma, and other neuromuscular disorders. One promising approach attempts to decipher motor imagery from the electroencephalogram (EEG) by recording sensorimotor rhythms (SMRs), which predictably change in response to real and imagined movements (Pfurtscheller and Neuper 2001; He et al. 2013). The intuitive and continuous nature of SMR-based BCIs enables the extension of the user through the control of virtual objects, drones, wheelchairs, and robotic arms (Wolpaw and McFarland 2004; Tsui et al. 2011; Lafleur et al. 2013; Edelman et al. 2019). Users, for example, might imagine turning a steering wheel with their left hand to direct a wheelchair to the left or drink independently with a robotic arm after imagining reaching for a cup.

However, major challenges remain in the translation of BCIs from the lab to the clinic— specifically, long training times and variable user proficiency currently prevent the widespread adoption of BCI technology. At least 20% of users are unable to control typical BCIs proficiently even after extensive training while performance fluctuations prohibit the safe use of robotic end effectors for daily activities (Blankertz et al. 2010). Though the vast majority of recent work has emphasized the development of decoding algorithms to improve BCI control, a promising complementary approach is to discover how individuals can be trained to interact with the system more effectively (Guger et al. 2000; Edelman et al. 2016; Perdikis et al. 2018). How individuals learn to control a BCI remains largely unknown; the paucity of research describing the evolution of explicit learning metrics—such as brain signal modulation—while the user gains control over the device remains a major hindrance to the development of effective training protocols (Perdikis et al. 2018).

Preliminary work has suggested mind-body awareness training (MBAT; e.g. yoga, meditation, etc.) may help to overcome some of the challenges BCI users face—as experienced practitioners demonstrate higher levels of BCI proficiency—however no research to date has generated a convincing explanation of the underlying mechanisms of this influence (Cassady et al. 2014; Tan et al. 2014).

Current theories of neurofeedback learning indicate that BCI control is achieved through the coordination of specific distributed brain networks (Ninaus et al. 2013; Sitaram et al. 2017), and these networks overlap significantly with those altered by meditation practice (Fox et al. 2014, 2016; Tang et al. 2015). While EEG does not allow the precise spatial localization of the activity in these networks, it is ideally suited to examine the electrical/oscillatory correlates of their communication.

Specifically, MBAT’s impact could result from meditation’s effects on alpha band power. Energy at these frequencies serves as the input to the decoder’s that control BCI (Schalk et al. 2004). Additionally, meditation has been shown to modulate specifically energy within this band. For example, expert meditators display greater alpha power during rest, a trait associated with greater BCI proficiency (Cahn and Polich 2006; Blankertz et al. 2010). Increases in alpha power have also been observed following the recognition of mind wandering, which mindfulness programs train individuals to monitor and may additionally reflect a learned ability to inhibit task irrelevant activity (Klimesch et al. 2007; Jensen and Mazaheri 2010; Braboszcz and Delorme 2011a; Mathewson et al. 2011; Roux and Uhlhaas 2014; Clayton et al. 2015; Rahl et al. 2017).

More generally, the emphasis on present-moment experience may allow expert meditators to self-regulate brain activity which could translate into enhanced BCI control (Faller et al. n.d.; Lutz et al. 2004; Garrison et al. 2013; van Lutterveld et al. 2017). Self-regulation supported by attentional control, emotional control, and self-awareness may additionally help users aim at a state of effortless relaxation, which has been hypothesized to improve BCI control (Witte et al. 2013; Tang et al. 2015).

While intuitive conjectures arise from research claiming MBAT enhances factors hypothesized to underlie BCI proficiency, to our knowledge none have been formally tested. We therefore designed and conducted a large-scale longitudinal BCI study to test whether and how short-term MBAT affects the learning and ultimate performance of SMR-based BCI control (Fig. 1A). A major goal of this work was to discover the neural bases of MBAT’s effects on SMR-based BCI performance. We predicted MBAT would confer this advantage through enhancing the capacity to: (1) sustain attention, (2) generate motor imagery, and/or (3) intentionally modulate rhythmic synchronization and desynchronization of neural activity during motor imagery or rest. This rigorous examination of how users learned to control the device and which aspects of this process could be accelerated or enhanced by MBAT should lead to better BCI training methods for larger populations of patients.

**Fig. 1.**
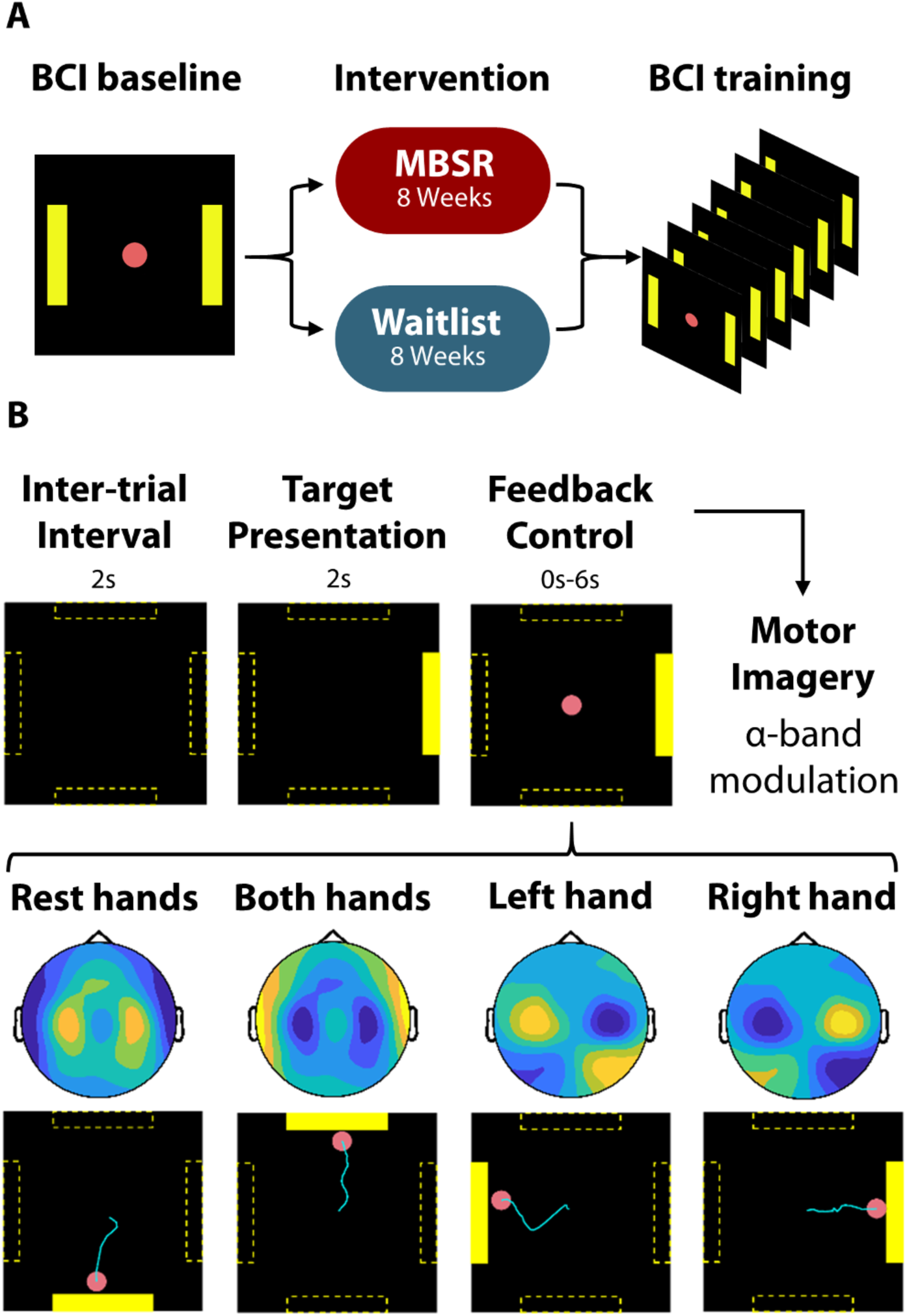
BCI training and control. **A**, Experimentation timeline for BCI training. Participants visited the lab for an initial assessment of BCI proficiency followed by random assignment to either the mindfulness-based stress reduction (MBSR) course or an 8-week waitlist condition. After the intervention, BCI learning was assessed through a series of up to 10 training sessions occurring 3 times per week. **B**, BCI task. Following a 2 s inter-trial interval, a target appeared on the screen for 2 s indicating where the cursor should be directed. During the feedback control period, the participant was given up to 6 s to direct a cursor toward the correct target. The cursor was controlled by changes in alpha power measured over bilateral motor cortices. During lateralized motor imagery, reductions in alpha power (blue in scalp images) are typically observed over the contralateral motor cortex, and weaker increases in alpha power (yellow in scalp topographies) are observed over the ipsilateral motor cortex. Bilateral motor imagery typically results in bilateral desynchronization of alpha activity relative to rest. In this way, alpha power asymmetry controlled lateral motion while the contrast between alpha power during motor imagery and rest controlled vertical motion.

## Materials and Methods

### Experiment Design

During the BCI tasks, users attempted to steer a virtual cursor from the center of the screen out to one of four targets (Fig. 1B). Participants initially received the following instructions: “Imagine your left (right) hand opening and closing to move the cursor left (right). Imagine both hands opening and closing to move the cursor up. Finally, to move the cursor down, voluntarily rest; in other words, clear your mind.” In separate blocks of trials, participants drove the cursor toward a target that required left/right (LR) movement only, up/down (UD) only, and combined 2D movement (2D). Cursor control was determined by the user’s ability to modulate the mu rhythm—an EEG oscillation in the alpha band (8-12 Hz) recorded over sensorimotor areas—such that alpha power asymmetry governed lateral motion while the contrast between alpha power during motor imagery and rest determined vertical motion (a standard BCI approach) (Schalk et al. 2004; He et al. 2013).

Following an initial baseline assessment of BCI performance, participants were randomly assigned to the MBSR group or a waitlist control group, respectively. The MBSR intervention seeks to teach mindfulness, defined by MBSR’s founder Jon Kabat-Zinn as “awareness that arises through paying attention, on purpose, in the present moment, non-judgmentally” (Kabat-Zinn 1982). Following the eight-week MBSR course, or comparable waiting period, participants returned to the lab for 6-10 sessions of BCI training. Groups (MBSR, n = 33, and Control, n = 29) were compared across various metrics of BCI performance and electrophysiological activity throughout training. After completion of the last session, the wait-list control subjects were provided with the opportunity to join an MBSR class comparable to that provided to the experimental subjects at a later date.

### Participants

All participants provided written informed consent and the study was approved by the Institutional Review Boards of the University of Minnesota and Carnegie Mellon University. In total, 144 participants were enrolled in the study and seventy-six participants completed all experimental requirements. Seventy-two participants were assigned to each intervention by block randomization, with 42 participants completing all sessions in the experimental group (MBSR before BCI training; MBSR subjects) and 34 completing experimentation in the control group. Only participants without (or with negligible) prior yoga or meditation experience were included in our study.

Typical participant involvement required roughly 6 hours/week over the course of 4 months, which we believe accounted for the high attrition rate. No significant differences were observed between groups (dropped vs completed / experimental vs control) in terms of demographics or initial BCI performance. A power analysis suggested a sample size of 40 participants in each group to achieve a statistical power of 0.9. The first group of participants (MBSR, n = 10; control, n = 11) completed 6 BCI sessions, however results indicated learning was still taking place; thus the experiment was extended to 10 BCI session.

Four subjects were excluded from the analysis due to non-compliance with the task demands and one was excluded due to experimenter error. Further, to evaluate the learning of BCI control, analysis focused on those that did not demonstrate ceiling performance in the baseline BCI assessment (accuracy above 90% in 1D control). The results describe data collected from 33 MBSR subjects (Age = 42 +/- 15, F = 26) and 29 controls (Age = 36 +/- 13, F = 23).

### MBSR

MBSR training consisted of a class that met weekly for 2.5 to 3 hours per class along with a 7.5 hour silent retreat during the sixth week of the class. Participants were also assigned home practice of 1 hour per day, 6 days per week. The content of the course was standard for MBSR, and included “body scans” and breath-focused meditation. Class attendance was recorded and participants were asked to use the Insight Timer (Insight Network, Inc., CA) to record their home practice. MBSR participants recorded on average 21.52 +/- 1.33 hours of formal practice outside of class throughout the course. Classes were taught by instructors of the University of Minnesota Earl E. Bakken Center for Spirituality & Healing.

### Measuring mindfulness

A validated behavioral measure of mindfulness, breath counting accuracy, was used to assess the effects of the MBSR intervention (Levinson et al. 2014). In separate 18 min sessions before and after the intervention, participants were asked to count their breaths in cycles of nine (inhale and exhale counting as one), pressing one button for the first eight breaths, and a second button for the ninth (Levinson et al. 2014). Accuracy was quantified as the number of correctly labeled breath cycles divided by the total number of cycles.

### BCI cursor control

BCI experiments were conducted using BCI2000, a general-purpose software platform for BCI research. Each BCI session consisted of 6 runs (25 trials each; ∼3 minute duration) of LR, UD, and 2D conditions for a total of 18 runs per day. As sessions progressed, participants were encouraged to find their own strategies. The control signal was extracted as different combinations of the autoregressive (AR) spectral amplitudes of the small Laplacian filtered electrodes C3 and C4 in a 3 Hz bin surrounding 12 Hz normalized to zero mean and unit variance. The magnitude of the cursor movement was determined by the normalized AR amplitude difference, updated every 40 ms. Horizontal motion was controlled by lateralized alpha power (C4 - C3) and vertical motion was controlled by up and down regulating total alpha power (C4 + C3). BCI Accuracy was quantified by a percent valid correct metric, calculated as the number of hits divided by the total number of non-timeout trials (a timeout occurred when the cursor did not contact a target within 6 s).

### EEG acquisition and pre-processing

Participants wore a 64-channel EEG cap, which was set up according to the international 10–10 system. EEG was acquired using SynAmps RT amplifiers and Neuroscan acquisition software (Compumedics Neuroscan, VA). The scalp-recorded EEG signals were digitized at 1000 Hz and filtered between 0.1 to 200 Hz with an additional notch filter at 60 Hz and stored for offline analysis conducted using custom Matlab software (R2018b, MathWorks Inc.) in conjunction with the EEGLAB (V. 14.1.1) and FieldTrip (revision 20190419) toolboxes (Delorme and Makeig 2004; Oostenveld et al. 2011).

EEG data were first bandpass filtered between 1 and 100 Hz and downsampled to 250 Hz. Noisy channels with high impedance were rejected by visual inspection and spherically interpolated. The data were re-referenced to a common average. Ocular artifacts were removed using independent component analysis (ICA) and a template matching procedure. Runs were broken into individual trials which included pre and post inter-trial intervals (2 s each), target presentation (2 s), and feedback control periods (0.04-6 s), however only trials with feedback durations longer than 1 s were used for spectral analysis (described below). Finally, trials with excessive EEG variance were rejected on the basis of visual inspection. On average, 84% [63 trials] +/- 5.3% [4 trials] (mean +/- SD) of the trials of each type (ranging between 41% [31 trials] and 100% [76 trials]) were retained for analysis during each session. The number of trials per type did not differ between groups.

### Power spectrum analysis

Complex Morlet wavelet convolution was used to estimate the power spectrum of the EEG signals. A family of wavelets was created with 30 frequencies log spaced between 4 and 60 Hz with cycles increasing from 3 cycles at 4 Hz to 10 cycles at 60 Hz. The alpha band power was extracted by averaging a 3 Hz bin centered at 12 Hz.

A Fisher discriminant score—calculated as the magnitude of the mean difference in alpha power between trial types normalized by the pooled standard deviation across trials—was used to assess differences in spectral power during BCI operation (Perdikis et al. 2018). In order to estimate the average strength of the control signal during a given session, the Fisher score was calculated based on the same combination of electrodes used to control the BCI (described above). To obtain topographical maps of alpha power discriminability between trial types, the Fisher score was calculated separately for each electrode. Event related synchronization (ERS) is typically observed during rest where populations of neurons synchronize their activity (which leads to increases in alpha power) while event related desynchronization (ERD) is typically observed when motor neurons are recruited during motor imagery (which leads to independent/desynchronized computations and reductions in alpha power) (Pfurtscheller and Lopes Da Silva 1999). Individual trial type event-related desynchronization (ERD) was estimated by normalizing the alpha power during BCI operation by the alpha power during the 1s preceding the trial. Finally, changes in topological discriminability and ERD were calculated by subtracting each subject’s baseline discriminability/ERD from the average of their final 3 BCI sessions to provide a stable estimate of neural activity during trained BCI operation.

### Source imaging

To image the underlying source of alpha activity, dynamic imaging of coherent sources (DICS) was used to identify the spatial source distributions (Gross et al. 2001). The leadfield matrix was calculated using the Colin27 MNI template and boundary element modeling (BEM) in the Fieldtrip toolbox using 3,898 equally distributed grid points (8mm uniform spacing) in the gray matter. DICS spatial filters were constructed from the leadfield matrix and a cross-spectral density matrix to maximize the activity of interest at each specific grid point while suppressing the contribution of all other grid points. The spatial distribution of power was obtained by multiplying the spatial filters and Fourier-transformed sensor level data. A source space Fisher score was calculated by independently estimating alpha power for each trial type then calculating the Fisher score (as above) at each grid point.

### Statistical Analysis

Linear mixed-effects models were fit using the lme4 package (1.1-21) in R (3.5.0) and p-values were computed with the lmerTest package (3.1-0), using Satterthwaite approximation for degrees of freedom(Bates et al. 2014; Kuznetsova et al. 2017). For between session comparisons of BCI performance, BCI accuracy was modeled over time with fixed effects of session (levels: 11), group (levels: MBSR, control), and task (levels: LR, UD, 2D). Random effects included within subject factors of session and task. The average BCI control signal over time was modeled with fixed effects of session, levels, and task (levels: LR, UD). Models were initially fit with three-way interactions of group, task, and session. Fixed effects structures of the mixed-effects models were reduced stepwise by excluding non-significant interaction terms/predictors and compared using ANOVA ratio tests until the respectively smaller model explained the data significantly worse than the larger model (significant X 2 -test) (Kuznetsova et al. 2017). Cubic polynomial contrasts were computed for the session factor, however only linear and quadratic terms (in addition to intercepts for task) were used for the random effects. Residual plots were examined for approximate normality and homoscedasticity. Assumptions of normality and homoscedasticity were clearly violated when attempting to fit the estimated BCI control signal with cubic contrasts (presumably due to the large differences in Fisher score between the LR and UD tasks [Fig. 3A,3B]) thus a two parameter box-cox transformation was applied to the dependent variable of Fisher score instead.

**Fig. 2.**
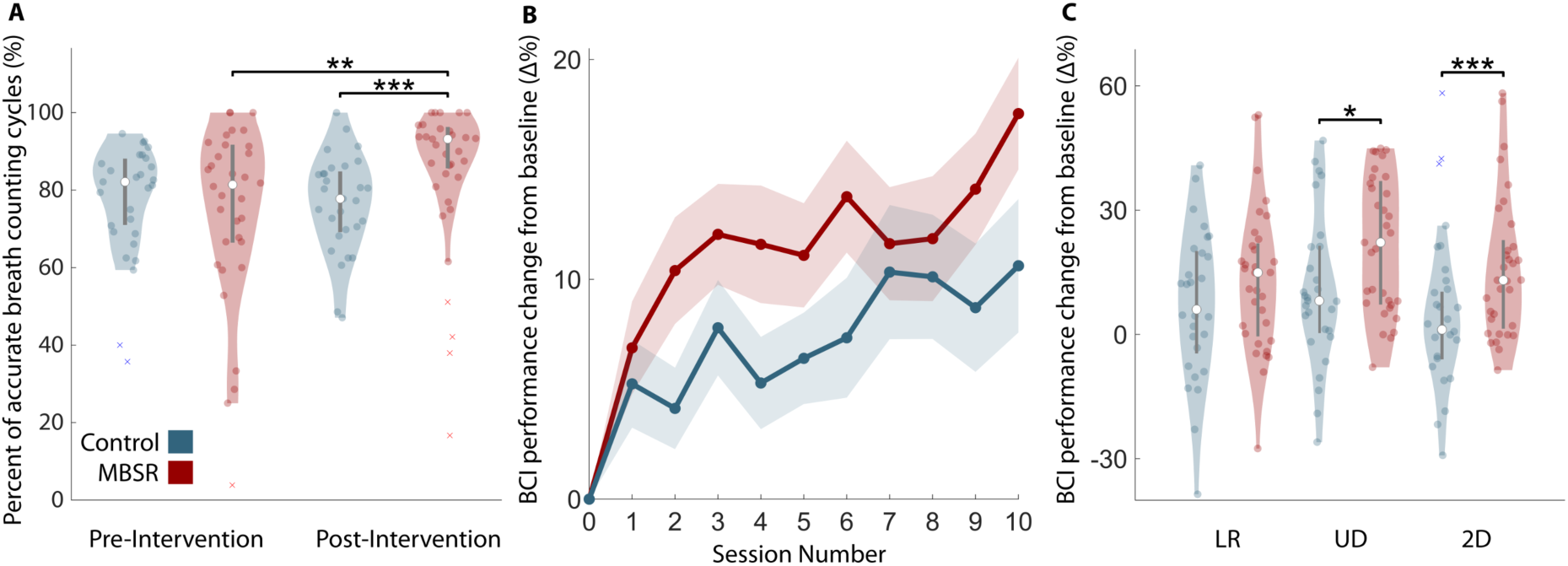
MBSR improves mindfulness and BCI control. **A**, MBSR improves breath counting accuracy. A significant interaction was observed between group and time suggesting MBSR affects breath counting accuracy (R-ANOVA P=0.029). Following the intervention, MBSR participants displayed significantly greater breath counting accuracies than their pre-intervention levels (WRS P=0.0027) and the post-intervention breath counting accuracies of controls (WRS P<0.001). **B**, The MBSR group shows greater improvements in average BCI performance throughout training. A significant interaction was observed between group and session number suggesting MBSR affects the learning of BCI control (mixed effects model, linear: P=0.003, quadratic: P=0.001, cubic: P=0.001). **C**, Significant improvements from baseline were observed in the UD (WRS, P=0.025) and 2D (WRS, P=0.002) tasks suggesting the observed interaction resulted from MBSR improving the learning of BCI control. Violin plots (**A**,**C**): Shaded area represents kernel density estimate of the data, white circles represent the median, and grey bars represent the interquartile range. Line plots (**B**): Shaded area represents +/- 1 standard error of the mean (S.E.M).

**Fig. 3.**
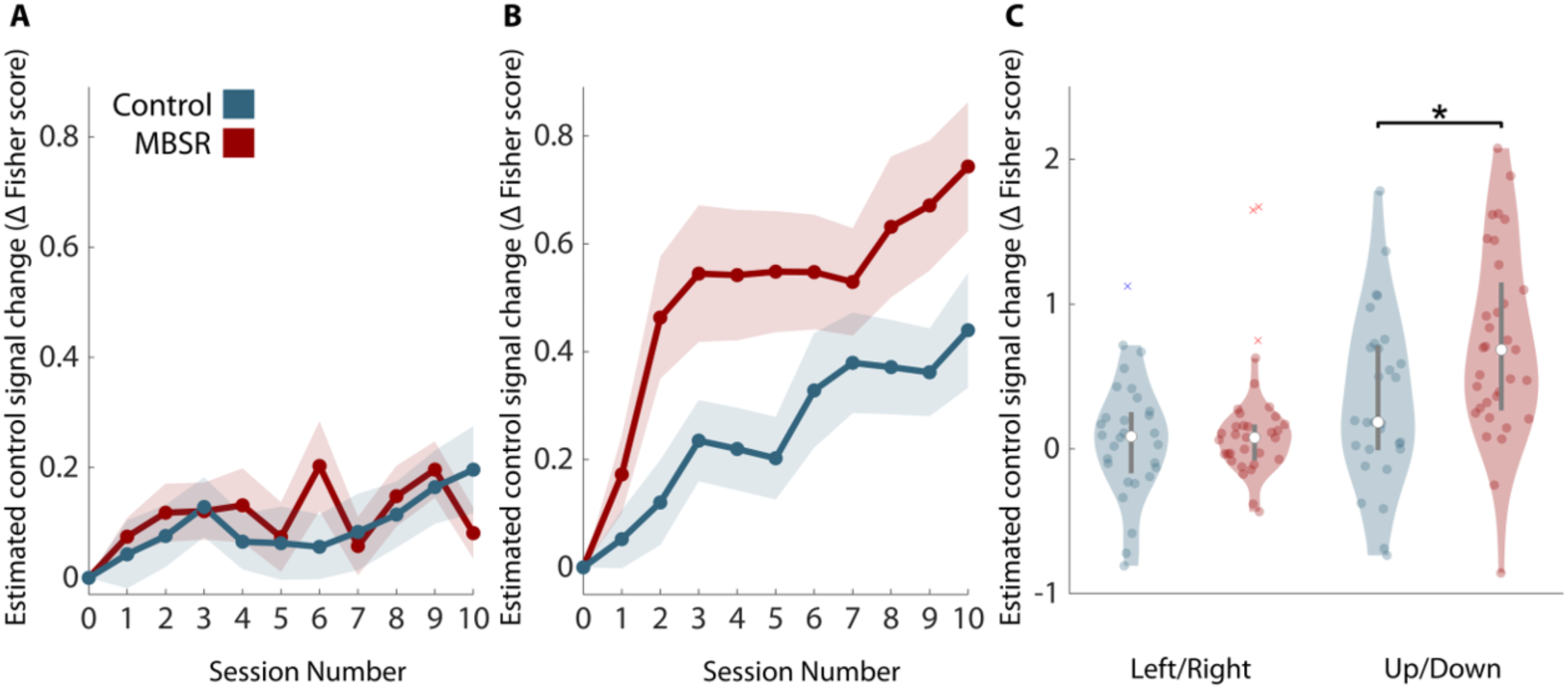
MBSR increases the contrast between motor imagery and rest. **A**, Change in estimated control signal modulation in the LR task during training. **B**, Change in control signal modulation in the UD task during training. A significant three-way interaction was observed between group, session number, and task suggesting MBSR changes how the control signal is modulated throughout training (mixed effects model, P=0.02). **C**, A significant difference in control signal modulation was observed in the UD (independent t test, P=0.019), but not the LR task, suggesting the role MBSR plays in BCI training is by changing the contrast between alpha power during motor imagery and rest. Line plots: Shaded area represents +/- 1 standard error of the mean (S.E.M). Violin plots: Shaded area represents kernel density estimate of the data, white circles represent the median, and grey bars represent the interquartile range.

A cluster-based permutation test (CBPT) statistic was used to assess the group differences in the changes of Fisher discriminant score recorded across the cortex during BCI training as well as for all source analysis (Oostenveld et al. 2011). A similar procedure was used for investigating the relationship between source space rest ERS and meditation experience, except that a t-transformation of the Pearson correlation was used as the statistical metric.

Repeated measures ANOVAs were used to test group by time interactions when data were found to be normally distributed using the Shapiro-Wilks test. The breath counting data displayed significant deviations from normality and was fit with a rank based analysis of linear models using the Rfit package (0.23.0) as a robust alternative to least squares (Kloke and McKean 2012). Significant findings were followed by independent post-hoc tests to aid inference. Non-parametric tests were used when group data was found to be non-Gaussian using the Shapiro-Wilks test. Confidence Intervals and Cohen’s D (difference in means/pooled standard deviation) were calculated when the parametric tests’ assumptions were met. All post-hoc inferences were two-sided at a Holm-Bonferroni corrected alpha level of 0.05 (0.025 per side). Outliers, defined as 3 scaled median absolute deviations away from the median, were removed for post-hoc statistical testing and are marked with an x in all plots. Source imaging was used as a post-hoc inference to supplement the sensor level results and 3 control subjects with outlying electrophysiological data were removed for these plots and tests.

## Results

### MBSR improves mindfulness and BCI control

The MBSR training was found to be effective, as indexed by breath-counting accuracy—a validated behavioral measure of mindfulness (Levinson et al. 2014). Using a rank based ANOVA (see Methods: Statistics), a significant interaction was found between group and time (*F*_1,60_= 4.92, P=0.029; Fig. 2A, Supplementary Table 1). Further analysis revealed that, after the intervention, the MBSR group breath counting accuracy was significantly greater than both their pre-intervention levels (Wilcoxon Rank-Sum test (WRS): Δ_med_=11.8%, IQR_MBSR-Pre_= 15.60%, IQR_MBSR-Post_=10.24%, Z_(n1 =32,n2 =29)_=3.00, P=0.0027) and the post-intervention levels of controls (WRS: Δ_med_=15.40%, IQR_MBSR-Post_= 10.24%, IQR_control-Post_=15.18%, Z_(29,29)_=4.14, P<0.001), suggesting MBSR participants were able to learn the techniques taught in the class and apply them in other contexts.

Critically, the MBSR group showed greater improvements in BCI performance than controls across training (Fig. 2B). This effect was captured by a significant interaction between the independent variables of session number and group (MBSR and control) in a linear mixed-effects model for the dependent variable of BCI accuracy (linear *t*_184_=3.06, P=0.003, quadratic *t*_1119_=-3.25, P=0.001, cubic *t*_1528_=3.26, P=0.001, see Supplementary Material: Statistical modeling; Supplementary Table 2; Supplementary Figure S1). The MBSR group displayed significantly greater improvements in overall BCI performance from baseline (Δ_µ_=8.98%, SD_MBSR_=14.55%, SD_control_=15.96%, independent *t*_60_=2.32, P=0.024, Cohen’s d=0.59) suggesting MBSR had a positive influence on BCI training.

To begin to probe *how* MBSR affected the BCI learning process, performance change in individual BCI tasks was compared between groups. The MBSR group displayed significantly greater improvements in accuracy within the UD (WRS: Δ_med_=14.06%, IQR_MBSR_= 29.35%, IQR_control_=20.61%, Z_(33,29)_=2.24, P=0.025) and 2D (WRS: Δ_med_=11.94, IQR_MBSR_= 10.24%, IQR_control_=15.18%, Z_(33,26)_=3.03, P=0.002) tasks compared to controls, however no significant differences were observed in the LR task (WRS: Δ_med_=8.89%, IQR_MBSR_=20.91%, IQR_control_=15.88%, Z_(33,29)_=1.06, P=0.29). While greater improvements in BCI accuracy from baseline were observed in the MBSR group, the proportion of participants reaching a proficiency criterion did not differ between groups (see Supplementary Material: Proficiency analysis).

### MBSR enhances the neural contrast between motor imagery and rest

We next examined the neural signals underlying BCI performance using the Fisher score which indexes the reliability of signal differences between conditions (e.g. up vs down in the UD task; see Methods). MBSR participants exhibited a greater ability to modulate their BCI-relevant EEG signals following training for the UD, but not the LR task. A significant three way interaction was observed between group (MBSR and Control), task (LR and UD), and session when a mixed-effects model was fit to the Fisher score over time (Fig. 3A,3B; *t*_1028_=2.33, P=0.02, see Supplementary Material: Statistical modeling; Supplementary Table 3; Supplementary Figure S2).

Examining the change in estimated control signals in the LR and UD tasks revealed that the differences in control signal modulation arose from alpha power contrast between motor imagery (up) and rest (down) (Fig 3C; Δ_*µ*_=0.38, SD_MBSR_=0.65, SD_control_=0.59, independent *t*_60_=2.42, P=0.018, Cohen’s d=0.62) rather than motor imagery of individual hands (Δ_*µ*_=0.017, SD_MBSR_=0.22, SD_control_=0.38, independent *t*_56_=0.22, P=0.83, Cohen’s d=0.058). Control analyses confirmed the difference in Fisher score was primarily driven by the difference in mean alpha power rather than difference in variance (see Supplementary Material: Control analysis—control signal components). These results were also confirmed by examining the UD and LR components of the 2D task, which served as a replication of our results; see Supplementary Material: 2D Analysis; Supplementary Figures S4-S6. Because significant differences in the LR control signal were not observed, subsequent analysis focused on the UD task exclusively.

Critically, MBSR-trained participants learned to dramatically alter both the amplitude and the spatial pattern of their alpha power during BCI control (Fig 4A). Cluster-based permutation tests (CBPT; see Methods: Statistics) revealed that changes in Fisher scores from baseline were significantly greater for MBSR participants than controls in the UD task across a wide range of electrodes including those over frontal, midline, posterior, and motor areas (Fig 4B; sensor space—cluster stat (maxsum(*t*_60_))=110.0, P=0.010; Fig 4C; source space—cluster stat (maxsum(*t*_57_)) = 6.67×10^3^, P=0.0001 see Supplementary Figure S3 for Fisher score evolution across time for LR vs UD comparison).

**Fig. 4.**
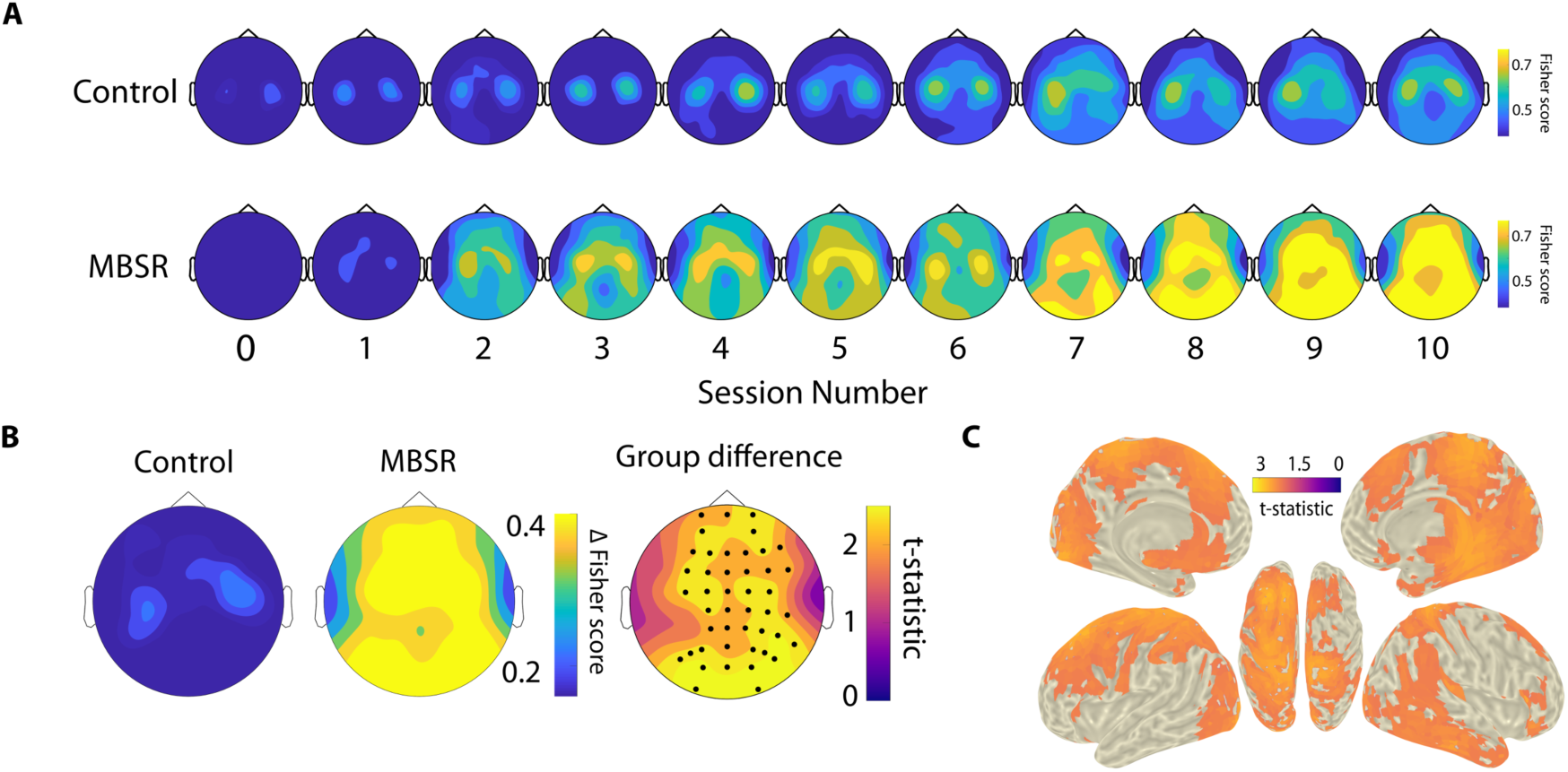
Alpha power contrast between motor imagery and rest is widespread. (A) The Fisher score was used to plot the difference between the distributions of alpha power during motor imagery vs rest at each electrode throughout training (x-axis = session number). During the UD task, the control group (top row) displayed the expected pattern in that the difference between motor imagery and rest is determined by the presence or absence of activity over the motor cortex. However, the MBSR group produced an entirely different pattern of contrast that evolves throughout training. Brighter colors represent greater differences in alpha power between trial types. **B**, Comparing the change in Fisher score across the cortex during UD control revealed the MBSR group learned to generate greater alpha power contrast between motor imagery and rest than controls across a wide range of electrodes (cluster-based permutation test, P = 0.01) **C**, Source imaging of the group difference in Fisher score change during the UD task again confirms the differences in learned alpha power modulation are widespread (cluster-based permutation test, P < 0.001).

### Mindfulness facilitates alpha power up-regulation during volitional rest

MBSR specifically led to increased alpha power up-regulation during rest trials. We separated trials into those controlled by motor imagery (up) and those controlled by resting (down) to determine which strategy contributed most to the observed results. The ability to modulate spectral power in the alpha band was calculated separately for each trial type by averaging alpha power during BCI operation and normalizing by the alpha power observed during the inter-trial interval to estimate event related desynchronization (ERD – reductions in alpha power) or event related synchronization (ERS – increases in alpha power). Power was averaged across the electrodes belonging to the significant cluster identified by the CBPT (Fig. 4B). A repeated measures ANOVA found a significant interaction between trial type and group when modeling change in ERS from baseline (*F*_1,60_=6.70, P=0.012; Fig. 5A, Supplementary Table S4). While no differences in alpha power modulation were found during motor imagery between MBSR subjects and controls (Δ_*µ*_=-0.014dB, SD_MBSR_=0.92dB, SD_control_=1.17dB, independent *t*_56_=-0.51, P=0.61, Cohen’s d=-0.14), a significant difference in the training-related increases in alpha power ERS was observed between groups during the rest trials (Δ_*µ*_=1.37 dB, SD_MBSR_=2.32dB, SD_control_=1.56dB, independent *t*_58_=2.61, P=0.011, Cohen’s d=0.68). Control analyses found no difference in passive resting state alpha power between the MBSR and control group indicating the reported differences in learned alpha ERS result from the MBSR subjects’ ability to *intentionally up-regulate* alpha power above and beyond the passive resting state during rest trials (see Supplementary Material: Control analysis—resting state power).

**Fig. 5.**
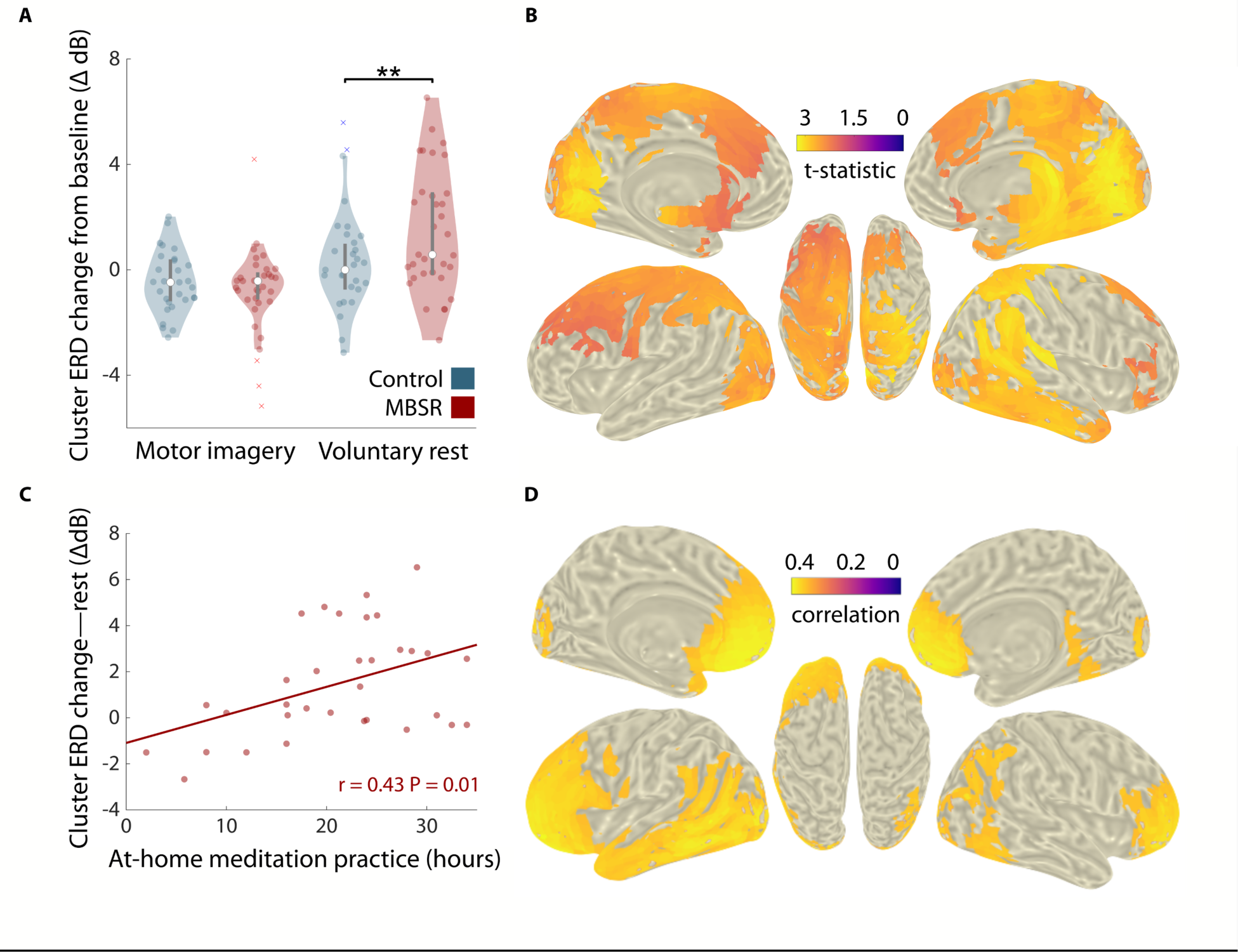
Localizing effects to rest. **A**, Averaging over the significant electrodes in Fig. 4B reveals the alpha power contrast between motor imagery and rest primarily results due to the learned ability to increase alpha ERS during rest trials (independent t-test P=0.011). **B**, The significant difference in the learned ability to synchronize alpha oscillations during rest covers roughly the same areas identified by the group difference in Fisher score in the source domain (cluster-based permutation test, P=0.0055). **C**, The training related increases in alpha ERS during rest displayed a dose dependent response with meditation practice suggesting more at home meditation practice led to greater changes in alpha ERS during BCI control (Pearson correlation, P = 0.012). **D**, Source imaging suggested meditation practice is most strongly associated with greater alpha ERS increases in frontal, temporal, and parietal areas (CBPT, P=0.028).

Source imaging revealed that the greatest difference between MBSR and control participants in the learned enhancement of alpha ERS during rest could be localized to roughly the same structures as the difference in Fisher score, suggesting the increase in alpha activity during rest was driving the improvement in BCI control (Fig. 5B; cluster stat(maxsum(*t*_57_))=5.52×10^3^, p=0.0055).

The learned ability to modulate alpha power during rest displayed a dose-dependent relationship with at home meditation practice and correlated with breath counting accuracy, suggesting both arise from a common MBSR ability. Further, this ability was found to predict final BCI performance indicating the contribution this skill plays in BCI control. We quantified the strength of association between alpha ERS within the identified cluster in Fig. 4B during BCI control and various behavioral variables. The change in alpha ERS power from baseline during rest trials was found to be significantly correlated with the hours of meditation practiced outside of the MBSR class (Fig. 5C; r_33_=0.43, P=0.012). Source imaging also suggested meditation practice is related to increases in alpha ERS with the strongest associations found in frontal, left temporal, and parietal areas (Fig. 5D; cluster stat(maxsum(*t*_33_))=2.29×10^3^, P=0.028). Finally, alpha ERS was found to correlate significantly with both ultimate BCI performance in the UD task (Fig. 6A; r_32_=0.51, P=0.0027) as well as post BCI training breath counting accuracy (Fig. 6B; r_31_=0.41, P=0.028). Collectively, these results suggest that the ability to up-regulate alpha power during volitional rest is related to mindfulness practice, results in better BCI control, and is related to the ability to pay sustained attention.

**Fig. 6.**
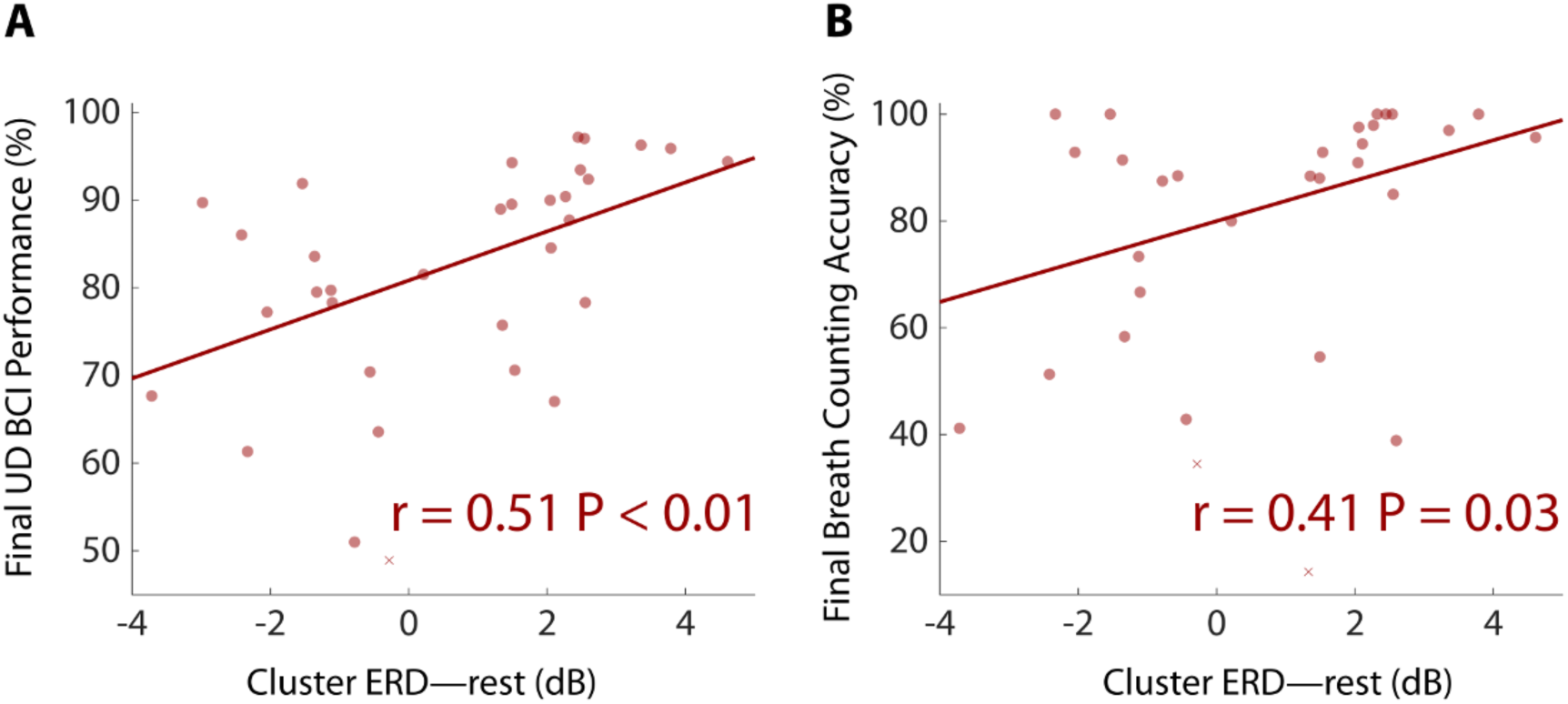
Rest trial alpha up-regulation predicts behavioral performance. Averaging over the significant electrodes in Fig. 4B demonstrates that alpha power up-regulation in the MBSR group correlates significantly with both **A**, ultimate UD BCI accuracy (Pearson correlation, P = 0.0027) and **B**, post BCI training breath counting accuracy (Pearson correlation, P = 0.028).

## Discussion

Our work demonstrates, in a large sample, that important aspects of BCI training can be enhanced through a behavioral intervention (Mcfarland and Wolpaw 2018; Perdikis et al. 2018). We identified the specific neural bases of the improvement in performance: Subjects trained in mindfulness were able to produce strikingly different patterns of alpha activity when the BCI task required volitional rest.

This ability to up-regulate this alpha band activity is functionally significant, as it has dose-dependent effects, predicts BCI performance, and is related to accuracy in a sustained attention task. Future work incorporating the identified mindfulness-based alpha pattern into the BCI decoder or combining mindfulness training with other interventions specifically targeting the sensorimotor rhythm could lead to even larger increases in accuracy, steeper learning curves, and more consistent performance for many different BCI training protocols.

We initially predicted MBAT would exert its effects through augmenting the ability to sustain attention, generate motor imagery, and/or influence brain rhythms; we found the strongest evidence for the latter. Increases in alpha band power have been commonly reported when comparing meditators and controls during both rest and meditation (Cahn and Polich 2006; Lee et al. 2018). Thus, prior research might predict the MBSR intervention simply taught participants to passively “rest better” by shifting baseline resting-state alpha power relative to controls (Blankertz et al. 2010). However, no differences in passive rest alpha power were observed between the groups after the intervention, nor throughout training. Rather, MBSR subjects demonstrated the ability to actively increase their alpha power while attempting to control the cursor during intentional rest.

This skill was not acquired directly from MBAT; that is differences were not evident in the first session after MBAT training, but instead were amplified throughout BCI training. Participants may have learned gradually to apply a skill they acquired from MBAT to help BCI performance, as this capacity was related to the amount of time invested in meditation practice outside of the class. This suggests that further practice could potentially lead to further gains in alpha ERS during BCI control (Cahn and Polich 2006; Hasenkamp et al. 2012; Saggar et al. 2012; Lee et al. 2018).

Our findings agree with prior work documenting experienced meditators’ ability to up-regulate alpha band power during meditation, however the nature of alpha oscillations during meditation remains a mystery (Cahn and Polich 2006; Brandmeyer and Delorme 2018; Lee et al. 2018). Research claiming that the recognition of mind wandering is associated with increases in alpha power and activation of the salience network (which in turn has been linked to alpha power) suggest alpha rhythms play an integral role in the “letting go” of spontaneous thought (Sadaghiani et al. 2010; Braboszcz and Delorme 2011b; Hasenkamp et al. 2012; Ellamil et al. 2016; Sadaghiani and Kleinschmidt 2016). In fact, reductions in neural noise, such as mind-wandering, are generally cited to explain why experienced mindfulness practitioners display enhanced abilities on tasks that probe attention (Lutz et al. 2009; Slagter et al. 2009; MacLean et al. 2010; Schoenberg et al. 2014; Colzato et al. 2015; Fox et al. 2016; Ziegler et al. 2019). The significant difference in the ability to follow the breath between the experimental and control groups subsequent to the MBSR intervention confirms previous reports that mindfulness is associated with reductions in mind wandering and the significant correlation between the ability to up-regulate alpha power during BCI control and breath counting accuracy suggests the MBSR participants may be engaging in meditative practices during the volitional rest trials of BCI control (Mrazek et al. 2012a; Levinson et al. 2014; Rahl et al. 2017).

The literature on alpha’s role in facilitating effective neural communication through the periodic inhibition of task irrelevant information agrees with the previous claim (Palva and Palva 2007; Jensen and Mazaheri 2010; Mathewson et al. 2011; Jensen et al. 2014; Sadaghiani and Kleinschmidt 2016; Miller et al. 2018; Chapeton et al. 2019). Our study found that the strongest relationship between meditation practice and increases in alpha power during BCI control could be localized to the prefrontal, left temporal, and parietal components of the DMN, which has been extensively linked to mind wandering (Eichele et al. 2008; Christoff et al. 2009; Hasenkamp et al. 2012; Fox et al. 2015; Ellamil et al. 2016). Therefore, the view of alpha oscillations as pulsed inhibition paired with significant improvements in sustained attention might indicate MBAT reduces mind wandering through stronger periodic gating of the DMN (Jensen and Mazaheri 2010; Mathewson et al. 2011; Brickwedde et al. 2019).

A limitation of the current work may be that MBAT does not improve BCI control in general. If the significant increases in breath counting accuracy represents improvements in sustained attention as we suggest, improvements across all domains of BCI control would be expected; however, no significant differences in electrophysiological activity were observed between the groups in the motor imagery task conditions. Alterations in the structure and function of sensorimotor cortex might require more meditation experience to develop (Fox et al. 2014, 2016). Alternatively, cuing the rest condition could promote mini-meditations leading to reduced mind wandering only during rest trials where transfer learning would be expected to be strongest (Mrazek et al. 2012b; Rahl et al. 2017).

The alpha increase that we observed was widespread across cortex. Alpha oscillations are traveling waves in the human neocortex, therefore the trial averaging employed in this analysis might obscure various metastable brain states that occur during BCI control (Zhang et al. 2018; Lozano-Soldevilla and VanRullen 2019; Roberts et al. 2019). In fact, a recent resting-state MEG study was able to decompose the DMN into anterior and posterior subcomponents which were active at different points in time (Vidaurre et al. 2018). Whether the increased in alpha power over the motor cortex results from radiating alpha sources or traveling waves within or between disparate metastable brain states is as yet unknown (Vidaurre et al. 2018; Zhang et al. 2018; Roberts et al. 2019). Regardless, since (1) most continuous BCI paradigms utilize rest as a state to be detected and (2) the decoder incorporates brain activity from both rest and motor imagery conditions in its decisions (i.e., there are not separate classifiers for rest and motor imagery), changing brain activity in one task improves classification of both conditions (evinced by the significant improvements in both 1D and 2D BCI control) (Perdikis et al. 2018; Edelman et al. 2019).

In conclusion, MBAT gives users of BCI the ability to learn to increase alpha power across the cortex on demand during rest. Future work detecting this mindfulness related alpha signal through techniques such as neural networks or common spatial patterns could lead to added improvements in BCI control as well as novel neurofeedback paradigms (Guger et al. 2000; Ang et al. 2008; Schirrmeister et al. 2017; van Lutterveld et al. 2017). Interventions or methods that specifically target sensorimotor rhythms such as noninvasive brain stimulation could additionally complement the performance gains reported in this study (Soekadar et al. 2015; Baxter et al. 2016; Kraus et al. 2016). Future work studying the ability of MBAT to promote the volitional up-regulation of alpha band power could further illuminate the role alpha oscillations play in neural communication and deepen our understanding of the neural correlates of attention, mind-wandering, and letting go of conscious thought.

## Acknowledgements

We would like to thank the following colleagues for their assistance in subject recruiting, data collection and analysis: Bhavani Sai Rohit Murakonda, Chris Coogan, Andy Huynh, Samantha Sherman, Alyssa Everson, Marisa Sanchez, Carter Ibister, James Kerber, Angeliki Beyko, Desirae Hammond, Kit Breshears, Taylor Boyle.

## Author contributions

B.H., S.E., J.R.S., and C.C.C designed the experiments. J.R.S. and C.C.C. collected the data. J.R.S. and H.J. analyzed the data. M.J.K. co-recruited subjects and helped facilitate the MBSR classes. J.R.S., S.E., C.C.C, and B.H. wrote the paper. All authors discussed the results and contributed to editing the manuscript.

